# Association between a functional polymorphism rs10830963 in *melatonin receptor 1B* and risk of gestational diabetes mellitus: an updated meta-analysis

**DOI:** 10.1101/141515

**Authors:** Yu Xiangyuan, Wang Qianqian, Qin Linyuan, Peng Lingxiang, Chen Zaiming, Qin Xiumei, Wang Yuchun, Shi Qingfeng, Yu Hongping

## Abstract

*The melatonin receptor 1B* (*MTNR1B*) as a candidate gene for gestational diabetes mellitus (GDM) on the basis of its association with T2DM, β-cells function and fasting plasma glucose (FPG) level. Many studies have investigated the association between *MTNR1B* polymorphism rs10830963 C>G and GDM risk, but the conclusion is inconsistent. PubMed, Google Scholar and CNKI databases were searched to identify eligible studies. Pooled OR with corresponding 95% CI was used to estimate the strength of the association between rs10830963 and GDM risk using a fixed- or random-effect model. 12 eligible studies with a number of 4,782 GDM patients and 5,605 controls were included in this meta-analysis. Results indicated that the variant G allele of rs10830963 polymorphism was significantly associated with an increased risk of GDM (CG vs. CC: OR=1.23, 95% *CI* = 1.12–1.34, *P*_heterogeneity_ = 0.23; GG vs. CC: OR=1.74, 95% *CI* =1.41–2.15, *P*_heterogeneity_ = 0.002). In the stratified analysis by ethnicity, similar results were found in Asians (CG vs. CC: OR=1.15, 95% *CI* = 1.04–1.28, *P*_heterogeneity_ = 0.74; GG VS. CC: OR=1.48, 95% *CI* =1.23–1.78, *P*_heterogeneity_ = 0.08) and in Caucasians (CG vs. CC: OR=1.49, 95% *CI* =1.25–1.77, P_heterogeneity_ = 0.28; GG vs. CC: OR=2.68, 95% *CI* =2.03–3.54, *P*_heterogeneity_ = 0.58).

## Introduction

Gestational diabetes mellitus (GDM) is defined as abnormal glucose tolerance with onset or first recognition during pregnancy_1_. Worldwide, it affects approximately 2~20% of all pregnancies. GDM has been shown to be associated with poor pregnancy outcome and substantial long-term adverse consequences for mothers and their offspring^3–6^. So far, the major risk factors related to GDM are older age at pregnancy, obesity, family history of T2DM and past history of GDM, previous poor obstetric history and genetics^7–13^. Insulin secretory defect accompanied by peripheral insulin resistance is an important characteristic of GDM^12^. Melatonin receptor 1B (MTNR1B) is an integral membrane protein that coupled to an inhibitory G protein and is expressed in pancreatic islets and pancreatic β-cells^13,14^. It has been found that the increased expression of MTNR1B on β-cells diminished intracellular cyclic cAMP levels, thereby inhibited the insulin secretion^15,16^. These findings suggested that MTNR1B may be involved in the development of GDM.

*MTNR1B* is located on human chromosome 11q21-q22, spanning about 22 kb and consisting of 3 exons and 1 intron. So far, 64 single nucleotide polymorphisms (SNPs) have been validated in the *MTNR1B* gene (http://www.ncbi.nlm.nih.gov/SNP), and some of which were reported to be associated with GDM risk. The SNP rs10830963 is located in the unique intron between exon 1 (+5.6 kb) and exon 2 (–5.9kb) of *MTNR1B* gene. Genotype-phenotype study of the SNP rs10830963 C>G showed that compared with the wild-type C allele of rs10830963, the variant G allele was associated with increased *MTNR1B* transcript levels in human islets^17^. Because of the functional consequence of rs10830963 C>G; many association studies have examined its effect on the risk of GDM^16,18–29^. However, these studies presented inconsistent results. To clarify the effect of the *MTNR1B* rs10830963 C>G on the risk of GDM, we therefore performed a meta-analysis with a total of 4,782 GDM patients and 5,605 controls from 12 published case-control studies.

## Materials and Methods

### Literature search strategy

We searched PubMed, Google Scholar and the Chinese National Knowledge Infrastructure (CNKI) databases for the association studies of *MTNR1B* rs10830963 polymorphism with the risk of GDM. The following key words were used: ‘MTNR1B’ or ‘Melatonin receptor 1B’, ‘gestational diabetes mellitus’ and ‘variation’ or ‘polymorphism’. All studies were published up to May 31, 2016. In addition, we manually searched for additional published studies on this topic in the references cited in the retrieved studies.

The included studies in this meta-analysis had to meet the following criteria: evaluation of the *MTNR1B* rs10830963 C>G polymorphism and GDM risk; case-control study; genotype or allele distribution information in cases and controls for calculating odds ratio (OR) with 95% confidence interval (CI); the study was written in English or Chinese. Accordingly, family-based studies, abstract, case reports, comments and reviews were excluded. If studies had overlapped subjects, only the largest study was included in the final analysis.

### Literature evaluation and data extraction

Two professional investigators (Yu XY and Wang QQ) independently reviewed the articles and extracted the data from all eligible publications. The following information was extracted from each included study: first author, year of publication, country, diagnostic criteria, genotyping methods, source of controls, number of cases and controls, genotype or allele distribution data of cases and controls, mean age, mean BMI and *P*-value of the Chi-square goodness of fit test for Hardy-Weinberg equilibrium (HWE) in controls.

### Statistical analysis

Deviation of genotype frequencies of the *MTNR1B* rs10830963 C>G polymorphism in controls from HWE was tested by using the Chi-square goodness of fit test, and a *P*-value less than 0.05 was considered a departure from HWE. The odds ratio (OR) and their 95% confidence interval (CI) were used to assess the strength of association between the *MTNR1B* rs10830963 C>G polymorphism and GDM risk. The heterogeneity across studies was assessed by the Q test, and was considered significant when a *P*-value less than 0.1^30^. A fixed-effect model was used to calculate the pooled OR if the heterogeneity was not significant, otherwise, the random-effect model was adopted^31^. The potential source of heterogeneity across the included studies was explored with meta-regression analyses. Stratified analyses were conducted by ethnicity (Asian and Caucasian). Both Begg’s and Egger’s tests were used to test for publication bias^32,33^. A *P*-value < 0.05 was considered as an indication for the potential presence of publication bias. Sensitivity analyses were done to assess the influence of individual study on the pooled ORs. All analyses were performed by using Stata software, version 12.0 (Stata Corp LP, College Station, TX, USA).

## Results

### Characteristics of include studies

The flowchart of study selection for this meta-analysis is presented in **Figure 1**. A total of 66 studies were found using our literature search strategy, of which 43 studies were excluded because of duplicates or not on the topic of polymorphisms and GDM risk. After full-text reviews of the remaining 23 articles, 11 studies were excluded for the following reasons: 2 studies were case only studies, 3 studies were review or meta-analysis articles, 6 studies didn’t focus on the topic of the *MTNR1B* rs10830963 C>G and GDM risk. Finally, 12 studies with 4,782 GDM patients and 5,605 controls were selected in this meta-analysis. Main characteristics of the included studies were shown in **Table 1**. The genotype frequency distributions of the rs10830963 C>G in controls were in agreement with HWE in all included studies except for the two by Vlassi^18^ *et al*. and Liu Q^34^ *et al*.

**Figure 1.**
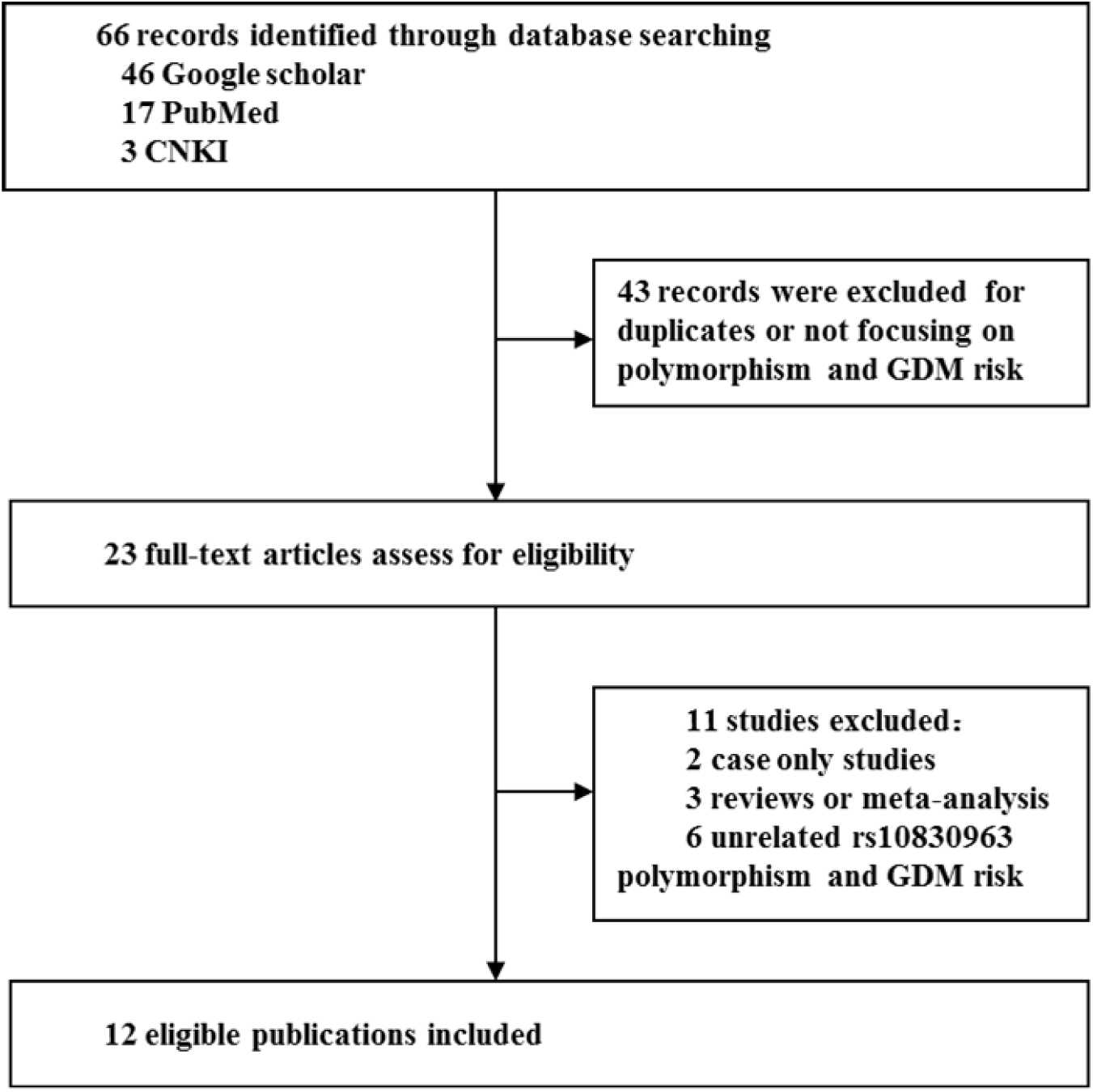
Flow chart of the process of identification of eligible studies

**Table 1.** Characteristics of the studies included in the meta-analysis

| Author, year<br>(reference) | Country | Diagnostic<br>criteria | Genotyping<br>methods | Controls | No. of<br>case/<br>control | Genotype distribution<br>(CC/CG/GG) | | MAF<br>case/control | Mean age<br>of cases/<br>controls | Mean BMI of<br>cases/<br>controls | $p_{HWE}$<br>for<br>controls |
| --- | --- | --- | --- | --- | --- | --- | --- | --- | --- | --- | --- |
|  |  |  |  |  |  | case | control |  |  |  |  |
| Deng Z,2011(28) | China | ADA | Sequencing | NGT | 87/91 | 23/38/26 | 31/45/15 | 0.52/0.41 | 31.8±4.6/29.7±3.5 | 23.6±3.0/21.5±2.4 | 0.84 |
| HuopioH,2013(29)* | Finland | ADA | Sequenom<br>Assay/TaqMan | NGT | 533/407 | 150/265/118 | 172/185/50 | 0.47/0.35 | 32.6/29.9 | 26.3±4.7/24.1±3.8 | 0.98 |
| Junior JP 2015 | Brazil | ADA | real-time PCR | Healthy | 183/183 | 102/61/20 | 113/66/4 | 0.28/0.20 | 32/29 | 32.0/25.4 | 0.11 |
| Kim JY,2011(30) | Korea | ADA | TaqMan | NGT | 908/966 | 217/435/256 | 294/469/203 | 0.52/0.45 | 33.1/32.2 | 23.3±4.0/21.4±2.9 | 0.53 |
| Kanthimathi S, 2015 | India | IADPSG | Sequenom<br>Assay | NGT | 518/910 | 155/259/104 | 309/437/164 | 0.45/0.42 | 28.1±5.0/26.3±5.5 | 27.5±2.4/23.3±4.7 | 0.66 |
| Li C,2013(31) | China | IADPSG | PCR-RFLP | NGT | 350/480 | 113/158/79 | 172/233/75 | 0.45/0.40 | 32.4±4.8/31.9±5.2 | 25.3±5.2/24.6±4.6 | 0.79 |
| Liu Q, 2015 | China | ADA | TaqMan | NGT | 674/674 | 162/334/178 | 195/362/117 | 0.51/0.44 | 31.6/32.1 | 24.4/25.2 | 0.02 |
| Qi J,2013(32) | China | IADPSG | Sequencing | NGT | 110/110 | 25/52/33 | 37/50/23 | 0.54/0.44 | 28.7±3.1/28.1±2.4 | NA/NA | 0.43 |
| Vlassi M,2012(33) | Greece | ADA | PCR-RFLP | NGT | 77/98 | 30/31/16 | 56/30/12 | 0.41/0.28 | 35.4±4.4/31.3±5.2 | 25.8±5.1/26.7±6.2 | 0.02 |
| Vejrazkova D,2014(35) | Czech | WHO<br>guideline | TaqMan | NGT | 458/422 | 169/227/62 | 206/184/32 | 0.38/0.29 | 34.1±6.1/34.8±15.1 | 24.3±4.9/23.7±4.2 | 0.48 |
| Wang Y,2011(34) | China | ADA | TaqMan | NGT | 700/1029 | 199/364/137 | 329/509/191 | 0.46/0.43 | 30.0/32.0 | 21.5/21.7 | 0.81 |
| Wang X, 2014 | China | ADA | PCR-RFLP | NGT | 184/235 | 62/89/33 | 69/121/45 | 0.42/0.45 | 28.2±3.8/27.9±4.1 | 21.2±1.8/20.7±1.4 | 0.53 |

ADA, American Diabetes Association; IADPSG, International Association of the Diabetes and Pregnancy Study Groups; NGT, Normal glucose tolerance; MAF, Minor Allele Frequency; BMI , Body Mass Index ;HWE, Hardy-Weinberg Equilibrium,.\* Genotype distribution data was calculated from allele frequency according to the law of HWE.

### Association between the *MTNR1B* rs10830963 C>G and GDM risk

As shown in **Table 2** and **Figure 2~3**, compared with the wild-type homozygous CC genotype, the variant CG heterozygous and GG homozygous genotypes was significantly associated with an increased risk of GDM (CG vs. CC: OR= 1.23, 95% *CI* = 1.12–1.34, *P*_heterogeneity_ = 0.23; GG vs. CC: OR=1.74, 95% *CI* = 1.41–2.15, *P*_heterogeneity_ = 0.002), respectively. An obvious heterogeneity across the studies was found in the homozygous genotype comparison.

**Figure 2.**
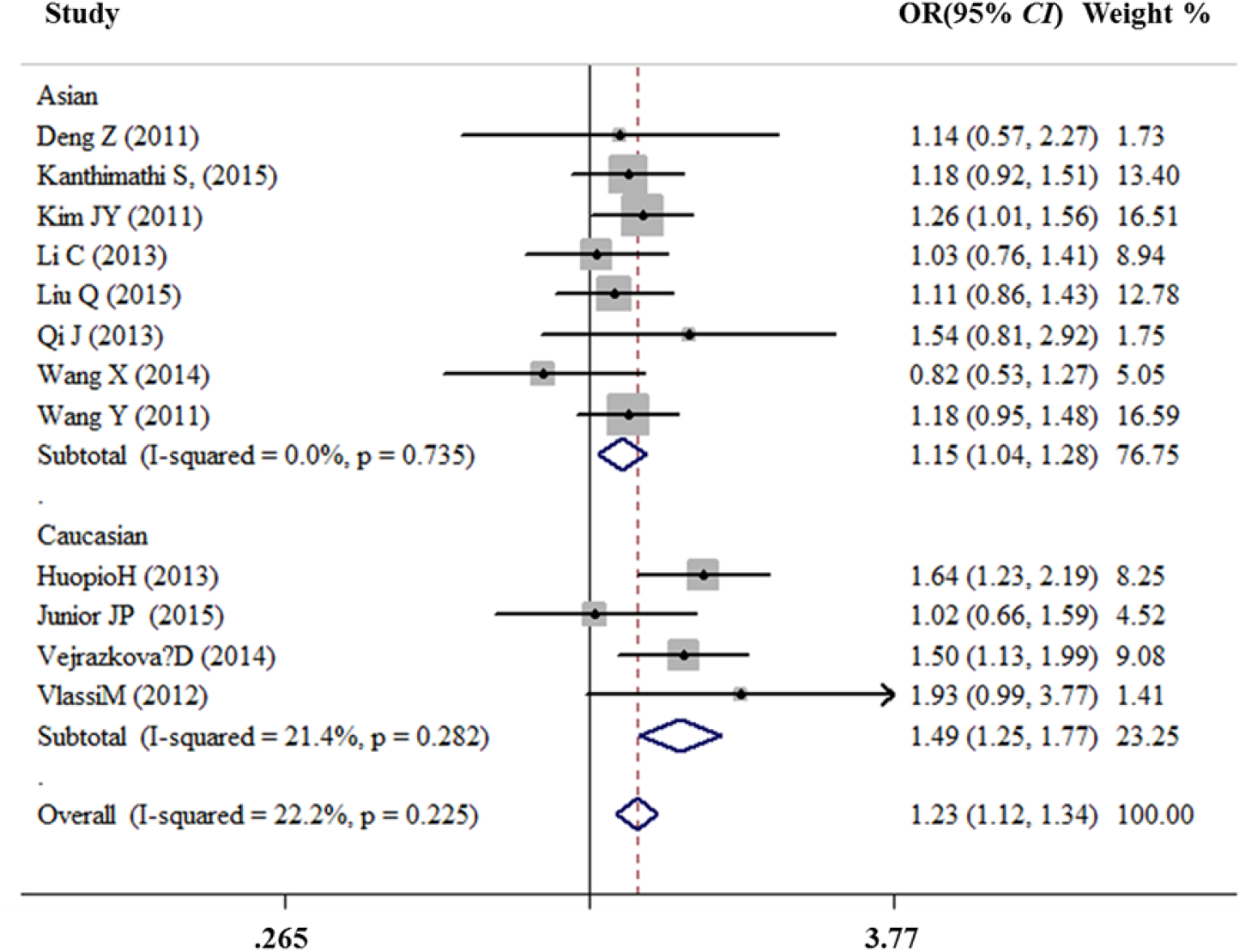
Forest plot risk of GDM associated with the MTNR1B rs10830963 (CG vs. CC)

**Figure 3.**
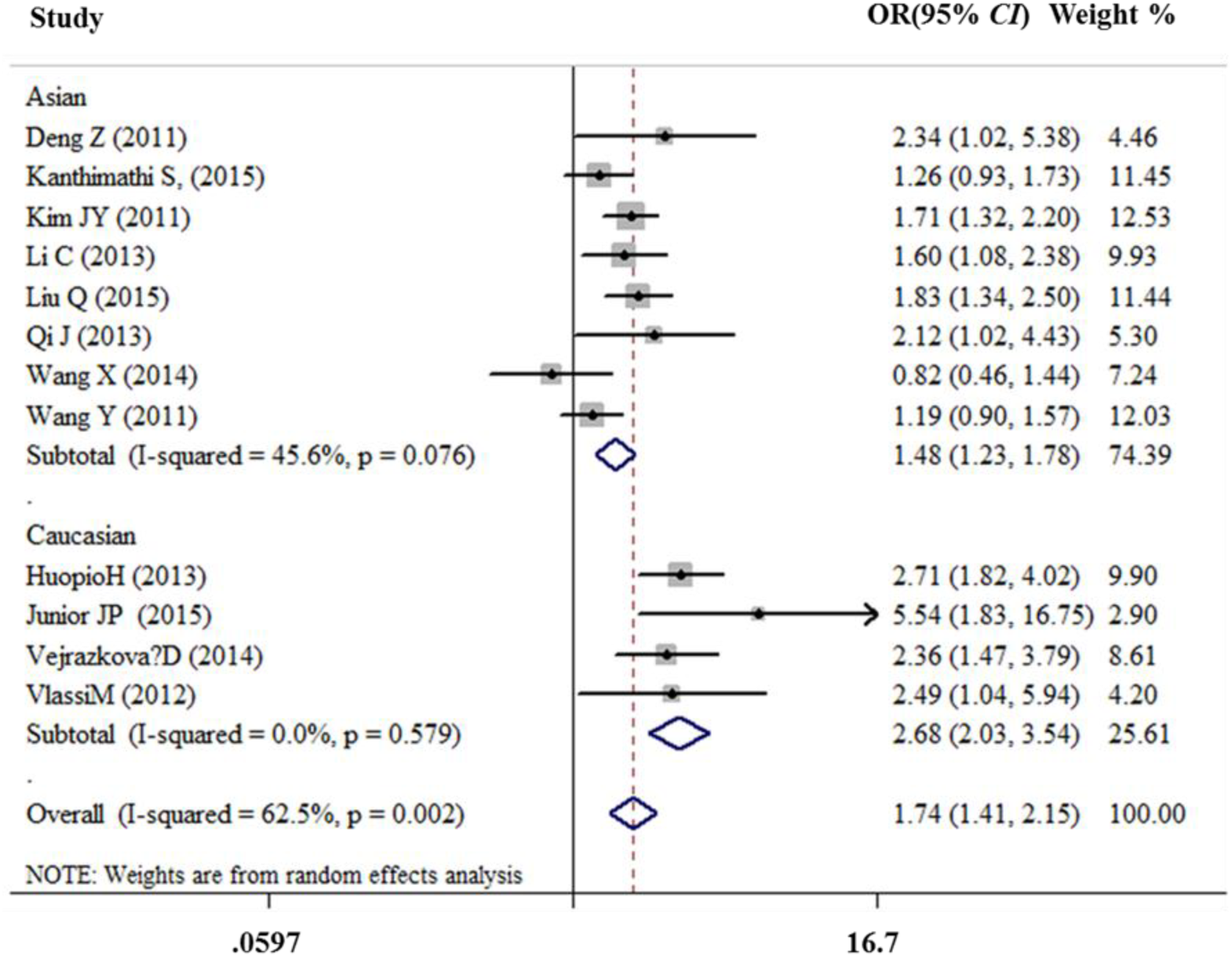
Forest plot risk of GDM associated with the *MTNR1B* rs10830963 (GG vs. CC)

**Table 2.** Meta-analysis of the *MTNR1B* rs10830963 polymorphism on GDM risk

| Subgroups | CG vs. CC |  | GG vs. CC |  |
| --- | --- | --- | --- | --- |
|  | OR(95% CI) | *p* Heterogeneity | OR(95% CI) | *p* Heterogeneity |
| Overall | 1.23(1.12-1.34) | 0.225 | 1.74(1.41-2.15) | 0.002 |
| Ethnicity |  |  |  |  |
| Asian | 1.15(1.04-1.28) | 0.735 | 1.48(1.23-1.78) | 0.076 |
| Caucasian | 1.49(1.25-1.77) | 0.282 | 2.68(2.03-3.54) | 0.579 |

In the stratified analysis by ethnicity, as shown in **Table 2** and **Figure 2~3**, compared with the homozygous CC genotype, the CG heterozygous genotype and the GG homozygous genotype were significantly associated with elevated risks of GDM in Asians and Caucasians (Asians: CG vs. CC: OR=1.15, 95% *CI* =1.04–1.28, Pheterogeneity = 0.74; GG vs. CC: OR=1.48, 95% *CI* = 1.23–1.78, *P*_heterogeneity_ = 0.08; Caucasians: CG vs. CC: OR=1.49, 95% *CI* =1.25–1.77, *P*_heterogeneity_ = 0.28; GG vs. CC: OR=2.68, 95% *CI* = 2.03–3.54, *P*_heterogeneity_ = 0.58). A modest heterogeneity across the studies was found in the homozygous genotype comparison in Asians. Subsequent leave-one-out analysis found that after removing the study by Wang Y19 *et al*. or Wang X^29^ *et al*., respectively, the heterogeneity among studies almost disappeared in Asian subgroup, suggesting that these two studies may contribute to the observed heterogeneity.

### Evaluation of heterogeneity

In this present study, the Q test was used to evaluate the heterogeneity across the included studies and the heterogeneity across studies was found in most of comparisons. We then used the meta-regression analysis to explore the source of heterogeneity and found that the ethnicity might contribute to the heterogeneity across studies in the overall analysis (CG vs. CC: t=–2.44, *P* = 0.035; GG vs. CC: t=3.17, *P* = 0.01).

### Sensitivity analyses

Sensitivity analyses of the association between the *MTNR1B* rs10830963 C>G polymorphism and GDM risk were performed to assess the stability of the pooled ORs under the CG vs.CC and GG vs.CC comparisons. The results showed that no single study dramatically influenced the pooled ORs (**Figure 4** and **Figure 5**).

**Figure 4.**
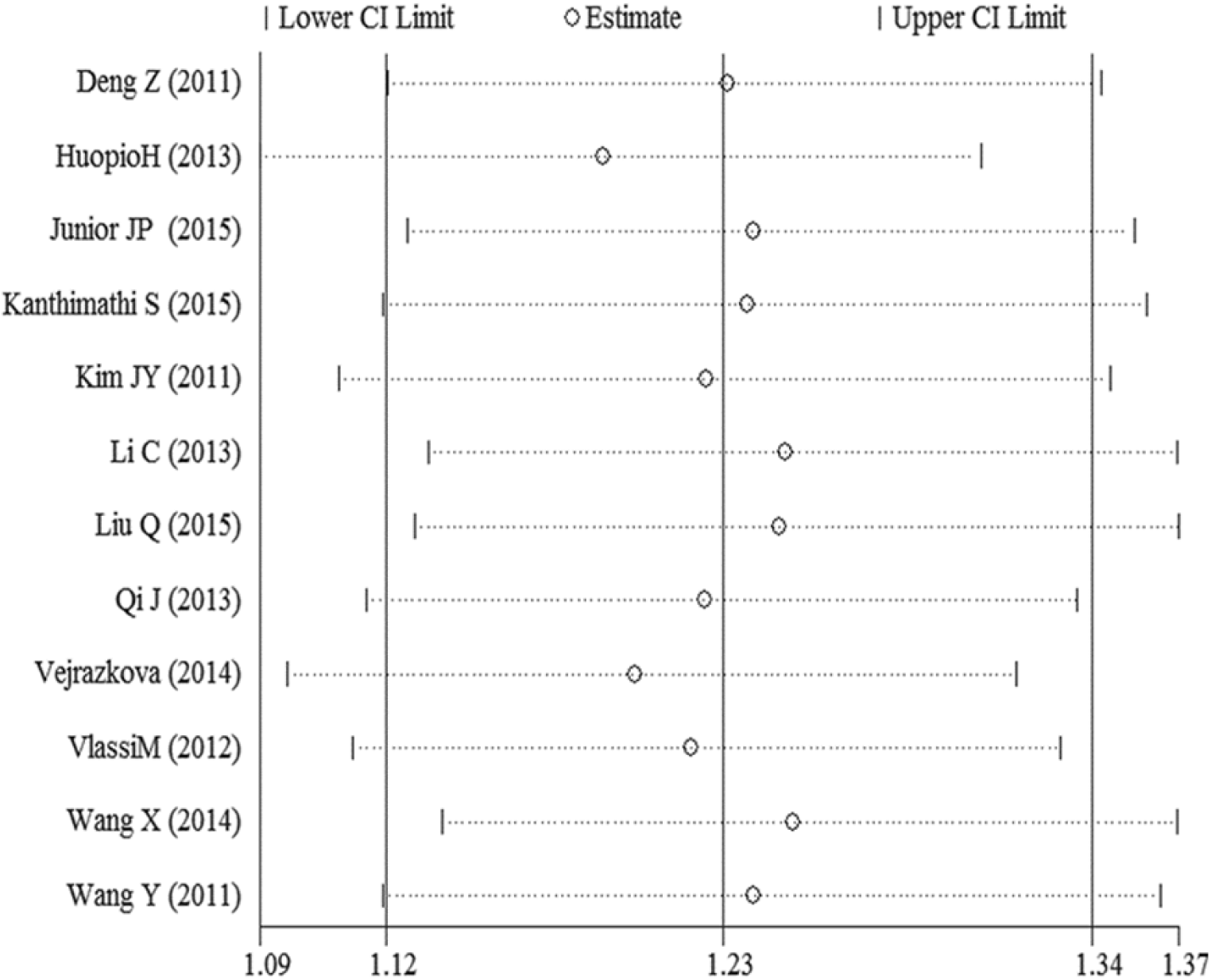
Sensitivity analyses of the association between the *MTNR1B* rs10830963 C>G GDM risk under the CG vs.CC comparison

**Figure 5.**
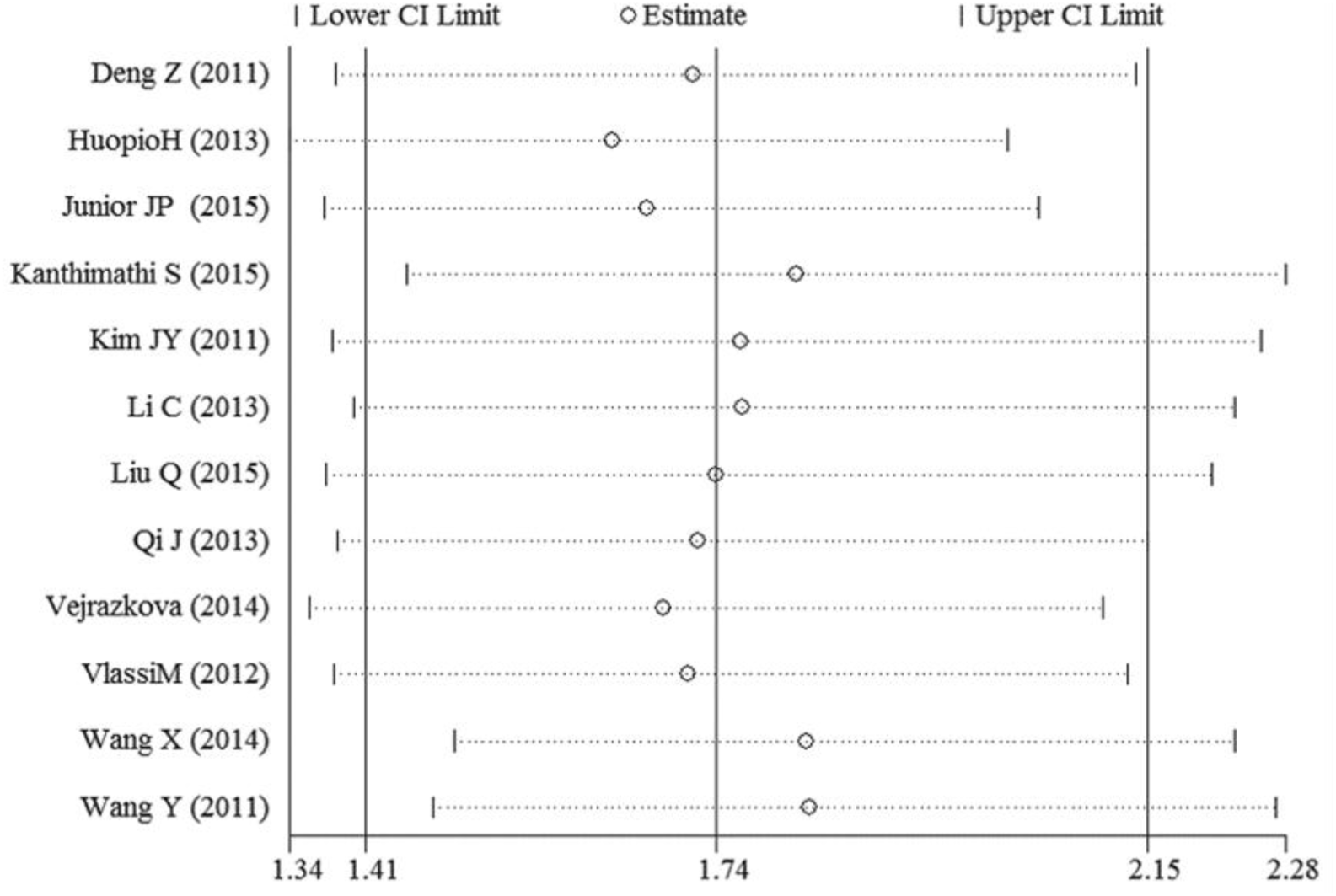
Sensitivity analyses of the association between the *MTNR1B* rs10830963 C>G GDM risk under the GG vs.CC comparison

### Publication bias

Both Begg’s and Egger’s tests were performed to evaluate the publication bias of the included studies. The shape of the funnel plots did not reveal any evidence of obvious asymmetry for all genetic models (**Figure 6** and **Figure 7**), and the Begg’s and Egger’s tests did not present any significantly statistical evidence of publication bias for any of the genetic models in the overall meta-analysis (CG vs. CC: *P_Begg’s_* = 1.000 and *P_Egger_’_s_* =0.905, GG vs. CC: *P_Begg’s_* =0.064 and *P_Egger’s_* =0.165). Neither funnel plots nor Begg’s and Egger’s tests detected any obvious evidence of publication bias in the subgroup analyses by ethnicity (data not shown).

**Figure 6.**
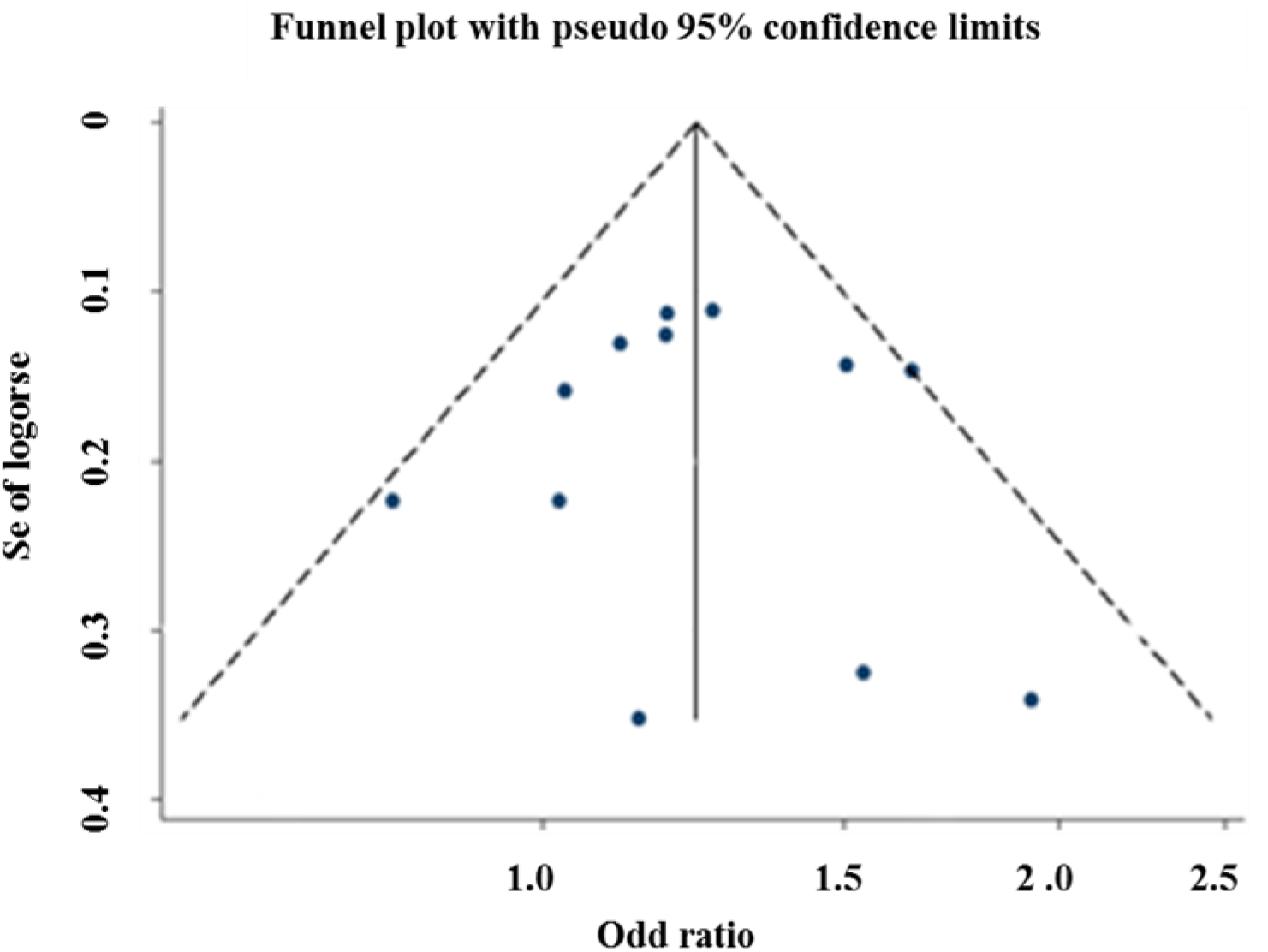
Begg’s funnel plot for publication bias test (CG vs. CC)

**Figure 7.**
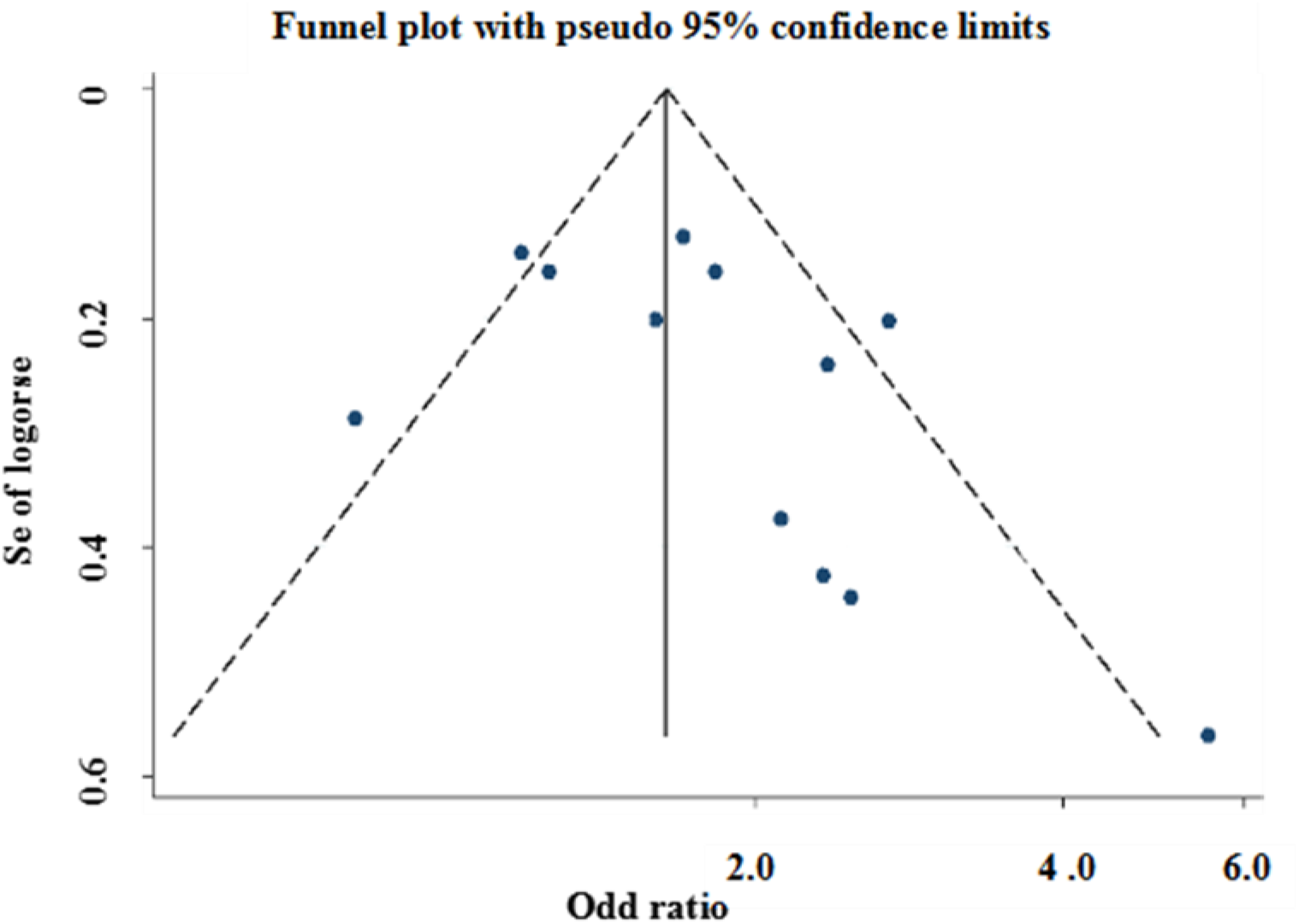
Begg’s funnel plot for publication bias test (GG vs. CC)

## Discussion

GDM could occur when the pancreatic islet β-cells were impaired by an increased insulin resistance during pregnancy^35^. MTNR1B is a G-protein coupled 7-transmembrane receptor and might influence pancreatic β-cells function and fasting plasma glucose (FPG) level^13,14^. Lyssenko^17^ *et al*.’s study showed that the increased expression of MTNR1B inhibited β-cells insulin secretion. A study of genetics and quantitative traits analysis of Palmer^36^ *et al*. revealed a significant association between *MTNR1B* and the glucose disposition index and acute insulin response amone T2DM. Meanwhile, experimental studies have confirmed that comparing with the wild-type C allele of rs10830963, the variant G allele caused an increased expression of *MTNR1B*^20–22,36–38^. To date, a number of case-control studies have been conducted to explore the effects of *MTNR1B* rs10830963 C>G polymorphism on the risk of GDM among different ethnic populations. However, the conclusions are still controversial. Huopio^23^ *et al*. found that the G allele of *MTNR1B* rs10830963 was significantly associated with GDM risk by increasing fasting plasma glucose and decreasing insulin secretion. However, Kanthimathi *et al*. reported that there was no significant association between rs10830963 polymorphism and the risk of GDM^24^. A previous meta-analysis by Zhang *et al*. observed a statistically significant association between the variant CG/GG genotype and GDM risk (OR=1.24, 95% *CI* =1.14–1.35)39. However, this meta-analysis only included 5 studies (4 studies for Asians and 1 for Caucasians) with 2,122 GDM patients and 2,664 control subjects, and was unable to do the subgroup analysis by ethnicity to reveal the effect of *MTNR1B* rs10830963 C>G polymorphism on GDM risk in different ethnic populations. Our updated meta-analysis summarized the evidence to date with 4,782 GDM patients and 5,605 controls and found that compared with the wild CC genotype, the variant CG and GG genotype were significantly associated with an increased risk of GDM, respectively. Furthermore, the subgroup analysis by ethnicity revealed that rs10830963 C>G polymorphism was significantly associated with the risk of GDM both in Asians and in Caucasians. Obvious heterogeneity across studies was observed in most comparisons in our meta-analysis. We then used the meta-regression analysis to explore the potential source of heterogeneity and found that the ethnicity might contribute to the observed heterogeneity. The current meta-analysis more comprehensively reflected the relationship between *MTNR1B* rs10830963 and GDM risk.

Some limitations should be point out in the current meta-analysis. First, although the Begg’s and Egger’s tests did not detect any significantly statistical evidence of publication bias, selection bias could exist because only published case-control studies were included. Second, the small sample size might limit the statistical power of the results in the subgroup analysis by ethnicity. For example, only four eligible studies with 1,251 GDM patients and 1,110 controls were included in the subgroup of Caucasian. Additional studies are warranted to further validate the effect of this functional polymorphism on the risk of GDM. Third, lack of individual-level data prevented us from making further analysis to identify any genotype-environment interaction between rs10830963 C>G and metabolic traits, such as FPG; pancreatic β-cell function, acute insulin response or indices for insulin sensitivity.

In summary, the current meta-analysis indicates that the variant G allele of *MTNR1B* rs10830963 C>G polymorphism may increase the risk of GDM. Given the limitations mentioned above, further large and well-designed studies of different ethnic populations are warranted to validate our findings.

## Additional information

### Acknowledgments

This study was supported by the Scientific Research Project of Education Department, Guangxi, China (grant No.KY2015YB223), the Science Research and Technology Development Project, Guangxi, China (grant No. 1412-4004-1-15).

### Competing financial interests

No competing financial interests exist.

